# MitoScribe single-cell molecular recorder logs graded signaling dynamics into mitochondrial DNA

**DOI:** 10.1101/2025.09.05.674553

**Authors:** Linhan Wang, Nikolaos Poulis, Deepak Srivastava, Seth L Shipman

## Abstract

Genetically encoded DNA recorders convert transient biological events into stable genomic mutations, offering a means to reconstruct past cellular states. However, current approaches to log historical events by modifying genomic DNA have limited capacity to record the magnitude of biological signals within individual cells. Here, we introduce MitoScribe, a mitochondrial DNA (mtDNA)-based recording platform that uses mtDNA base editors (DdCBEs) to write graded biological signals into mtDNA as neutral, single-nucleotide substitutions at a defined site. Taking advantage of the hundreds to thousands of mitochondrial genome copies per cell, we demonstrate MitoScribe enables reproducible, highly sensitive, non-destructive, durable, and high-throughput measurements of molecular signals, including hypoxia, NF-κB activity, BMP and Wnt signaling. We show multiple modes of operation, including multiplexed recordings of two independent signals, and coincidence detection of temporally overlapping signals. Coupling MitoScribe with single-cell RNA sequencing and mitochondrial transcript enrichment, we further reconstruct signaling dynamics at the single-cell transcriptome level. Applying this approach during the directed differentiation of human induced pluripotent stem cells (iPSCs) toward mesoderm, we show that early heterogeneity in response to a differentiation cue predicts the later cell state. Together, MitoScribe provides a scalable platform for high-resolution molecular recording in complex cellular contexts.

## INTRODUCTION

Cells integrate diverse and transient signals to guide development, maintain homeostasis, and adapt to environmental changes. While many of these signals are short-lived, their consequences can be long-lasting(*1, 2*). Thus, a full understanding of cellular systems requires information about both current cell states and as well as the molecular events that led to those states.

Conventional methods for probing cellular states—such as RNA sequencing and fluorescence imaging—offer powerful tools for molecular analysis but remain limited in key ways. Destructive techniques like RNA-seq capture only a snapshot of gene expression at a single time point, making it difficult to infer past signaling events. Live imaging of fluorescent reporters allows dynamic tracking but is restricted to optically accessible systems and constrained in multiplexing capacity(*3–7*). These limitations make it challenging to continuously monitor the temporal progression of molecular signals.

To address these limitations, recent efforts have explored ways of converting transient signals into stable genomic records, an approach known generally as molecular recording. Molecular recording platforms have been developed that use site-specific recombinases (SSRs)(*8–12*), CRISPR-based lineage tracing(*13–18*), inducible base editing(*19–22*), prime editing(*23–25*), or integrase-mediated barcoding(*26–31*) to write biological information into the nuclear genome. These approaches typically rely on engineered systems that introduce mutations into the nuclear genome in response to specific stimuli. However, the diploid nature of nuclear DNA inherently limits the dynamic range of such recordings at the single-cell level, where outcomes are constrained to a small number of discrete editing states: (1) unmodified, (2) heterozygously modified, or (3) homozygously modified. In addition, variability in editing efficiency and genomic accessibility further restricts the temporal resolution and information density of these systems.

Mitochondrial DNA (mtDNA) presents an alternative genomic substrate with unique advantages for molecular memory. Each mammalian cell contains hundreds to thousands of mitochondrial genomes, which are genetically isolated from the nucleus(*32, 33*). The recent development of bacterial DddA-derived cytosine base editors (DdCBEs) has made it possible to install targeted point mutations in mtDNA without introducing double-strand breaks(*34*). Using these mitochondrial base editors, we sought to leverage the copy number of mtDNA for encoding analog information, such as the strength or duration of a signal, in a reproducible and interpretable manner.

Here, we describe MitoScribe, a strategy for writing graded biological signals directly into the mitochondrial genome as neutral mutations using DdCBE mitochondrial base editors. These editors use a bacterial cytidine deaminase that is split into two halves. Each half is fused to a mitochondrial localization signal as well as an uracil glycosylase inhibitor (UGI) and a transcription activator-like effector (TALE) domain that targets the two halves of the deaminase to adjacent regions on the mitochondrial genome. We design these TALEs to reconstitute the deaminase over a cytosine in the mtDNA, which makes a silent mutation after the base is converted to thymine(*34*). The genes encoding the deaminase halves, which reside in the nucleus, are driven by promoters that are responsive to cellular signals that we wish to record (Fig. 1a,b). Critically, there are 100s-1,000s of copies of the mitochondrial genome in every cell. Therefore, rather than a signal leading to an allor-none outcome as in a Cre reporter system(*35*), in our MitoScribe system, these edits accumulate in proportion to pathway activity, that is, a weak signal leads to a low percentage of mitochondrial genomes being edited and a strong signal leads to a high percentage being edited (Fig. 1c). By coupling DdCBE activity to cellular pathways such as hypoxia, NF-κB, BMP and Wnt signaling, we generate single-nucleotide edits in mtDNA that reflect both the intensity and temporal extent of signal activation. We further show that these mitochondrial edits can be recovered alongside nuclear gene expression profiles in single cells through transcriptome sequencing with targeted enrichment of mitochondrial transcripts. Applying this approach to early mesodermal differentiation of human iPSCs, we uncover how historical signal dynamics shape cell fate decisions. Together, our findings highlight mtDNA as a genetically stable, high-capacity medium for recording biological events at single-cell resolution.

**Figure 1.**
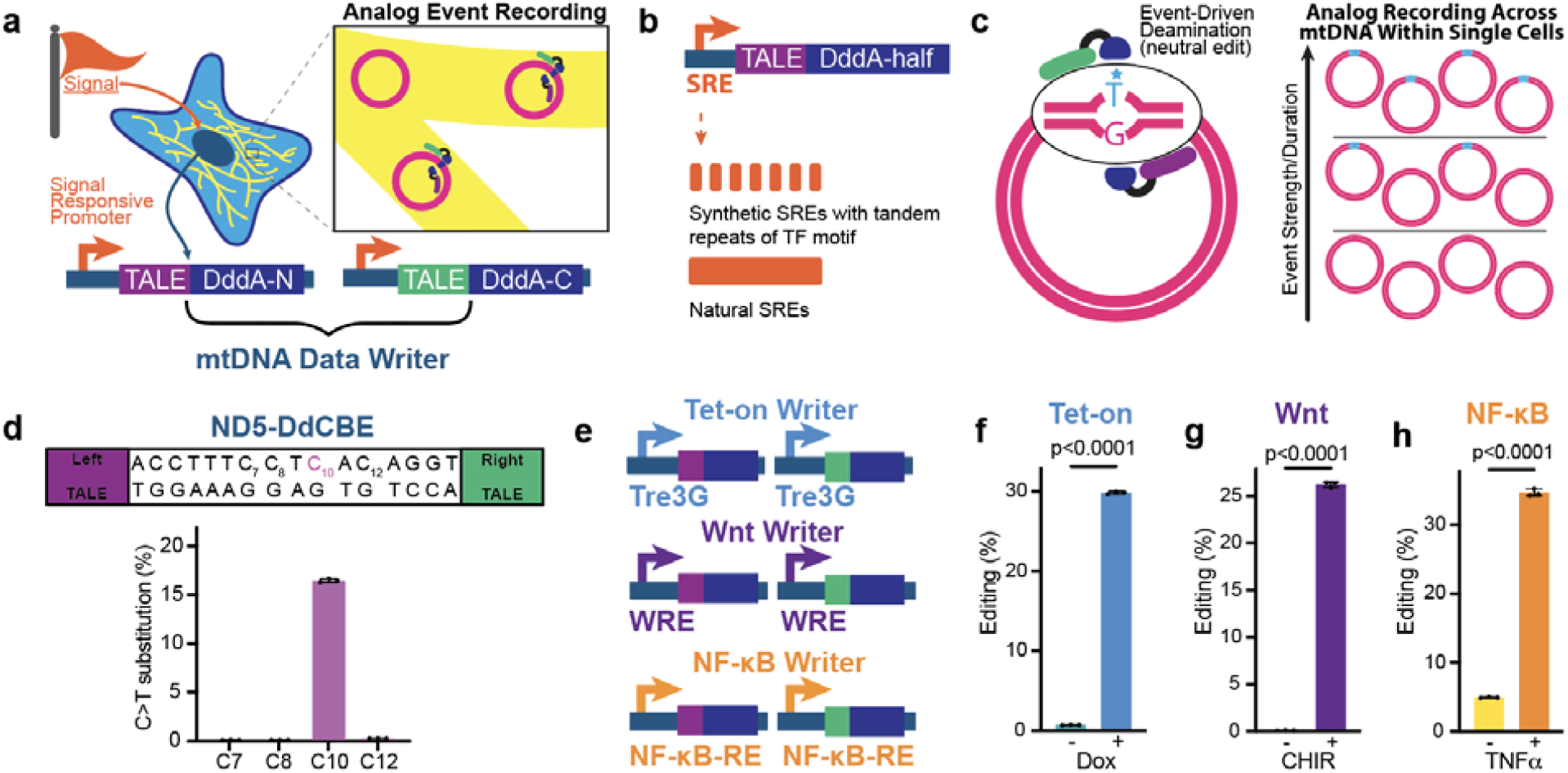
MitoScribe enables cellular events recording into mitochondrial DNA. **a,** Schematic of MitoScribe elements to record a signaling event. Two halves of a DdCBE base editor are placed under signal responsive promoters and targeted to adjacent sites on the mitochondrial genome via TALE domains. **b,** Synthetic signal responsive elements (SREs) with tandem motif repeats or natural SREs drive the expression of DdCBE halves. **c**, The event is written as a stable and neutral C to T transition distributed across mtDNA molecules within each single cell. The proportion of mtDNA with the edit indicates the integrated activity of the signal. **d**, The intended edit of the ND5-DdCBE and the frequency of C>T substitutions for the intended and bystander cytosines. Circles and error bars show the mean ± s.d. of 3 independent biological replicates. **e**, Schematic of three MitoScribe writers responsive to Tet-On (TRE3G), Wnt (TCF-LEF responsive element, WRE), and NF-κB signaling (NF-κB responsive element, NF-κB-RE). **f-h**, Signal responsive recording in cells transfected with each writer show editing upon pathway activation using 1 µg/ml doxycycline (**f**), 5 µM CHIR99021 (**g**), and 10 ng/ml TNFα (**h**). Bars show the mean ± s.d. of 3 independent biological replicates. P < 0.0001 by Student’s unpaired two-tailed t-test.

## RESULTS

### Cellular event recording into mtDNA via MitoScribe

To enable MitoScribe recording of cellular events into the mitochondrial genome without disrupting its function, we first identified a DdCBE base editor capable of precisely introducing a synonymous mutation in the *MT-*ND5 gene when constitutively expressed in human embryonic kidney 293T (HEK293T) cells for 2 days (Fig.1d and fig. S1a). Precision of the edit is critical to preserve mitochondrial function, which required empirical testing of TALE combinations, as it is not yet possible to predict whether a given editor will introduce non-synonymous and bystander mutations to nearby cytosines. For example, we tested a different TALE-DdCBE combination targeting the *MT-ND6* gene, which introduced multiple missense and nonsense mutations that result in partial loss of the gene product (fig. S1b). Using the *MT-*ND5 editor, neutral edits increased over 5 days while the plasmid was present in the cells, up to 35.9% as assessed by amplicon sequencing, and were retained through many cell generations and multiple passages when rechecked 15 days post transfection (fig. S1c).

We next placed the editor components under the control of signal-responsive promoters to create MitoScribe data writers, with the two halves of the deaminase carried on separate plasmids. We initially constructed a Tet-on writer, a Wnt writer, and an NF-κB writer (Fig. 1e). Then, we transiently co-transfected these MitoScribe writers into HEK293T cells and tested whether pathway activation led to editing using signal agonists: 1 µg/ml doxycycline (Dox)(*36*); 5 µM CHIR99021 (CHIR)(*37*); and 10 ng/ml TNFl⍰(*38*), respectively. In each case, we observed low background editing in the absence of the signal agonist and substantial editing in the presence of the agonist, with an overall editing efficiency of 29.8% for Tet-on recording (Fig. 1f), 26.2% for Wnt signaling recording (Fig. 1g), and 34.7% for NF-κB recording (Fig. 1h) 3 days post transfection. These results show that the MitoScribe approach can be used to record signal activation in mammalian cells.

### Recording of historical hypoxia signals

To assess whether MitoScribe could be used to record a cell’s exposure to changes in environment, rather than the addition of a chemical to the media, we focused on hypoxia, a physiologically relevant signal with well-characterized dynamics. We constructed a hypoxia-responsive MitoScribe writer under control of hypoxia response elements (HREs)(*39, 40*), enabling its expression in low-oxygen conditions, and transiently co-transfected the components into HEK293T cells. We observed production of the hypoxia MitoScribe writer protein after 2 days of exposure to low oxygen conditions (2% O_2_), but not in normoxic conditions (21% O_2_), confirming oxygen-dependent activation of the system (Fig. 2a). This environmentally responsive activation is recorded as a neutral edit to the mitochondrial genome (Fig. 2b). Neither the hypoxia or, crucially, the expression of the recorder led to a significant change in mitochondrial DNA (mtDNA) copy number (fig. S2a). With sustained hypoxic exposure, cells harboring the hypoxia writer showed progressive accumulation of the recording edit up to ∼10%, consistent with integration of signal duration, with substantially less accumulation in normoxia (Fig. 2c). As a control, we tested cells containing constitutively-expressed MitoScribe writers, which accumulated similar levels of edits in hypoxia or normoxia, up to 35.9% and 37.3%, respectively (fig. S2b).

**Figure 2.**
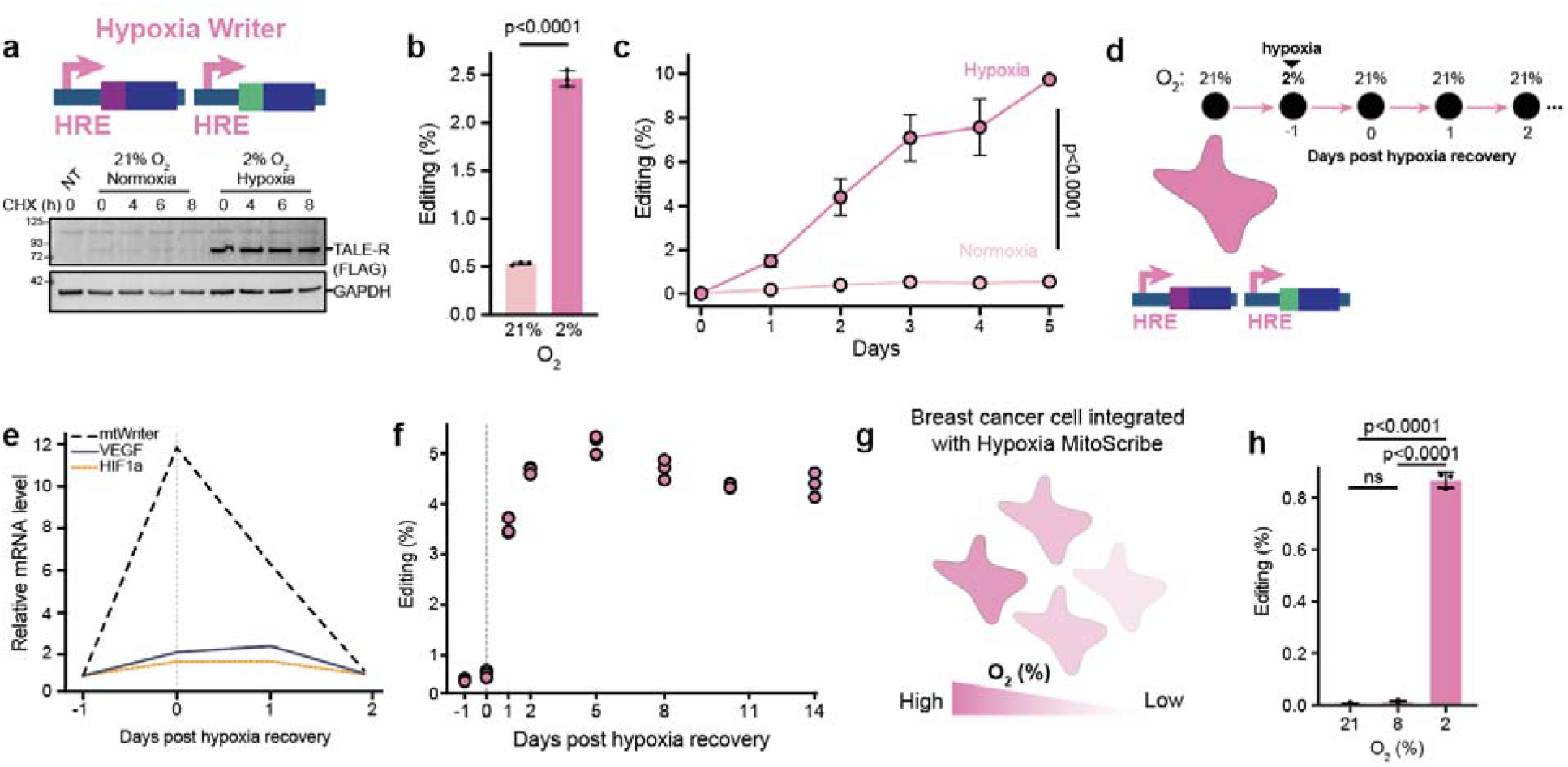
Recording and storing historical hypoxia signals with MitoScribe. **a**, Hypoxia-responsive MitoScribe writer is activated by low oxygen. Western blot of the hypoxia writer, showing production and stability of FLAG-tagged writer right half after 48 h in normoxia or hypoxia followed by treatment with 100 µg/ml cycloheximide (CHX). GAPDH was used as a loading control. **b**, Hypoxic exposure is recorded in the 2% of low oxygen condition. Circles and error bars show the mean ± s.d. of 3 independent biological replicates. P < 0.0001 by Student’s unpaired two-tailed t-test. **c**, Time-course experiment showing progressive accumulation of editing in HEK293T cells harboring the hypoxia MitoScribe writer during 5 days of hypoxia, but minimal editing over the same time-course in normoxia. Circles and error bars show the mean ± s.e.m. of 3 independent biological replicates. P < 0.0001 by unpaired two-tailed t-test**. d**, Schematic of transient hypoxia exposure for 24 h and normoxia recovery. **e**, Transcriptional response to transient hypoxia for hypoxia writer, VEGF, and HIF1α by RT-qPCR is over after 2 days of normoxia recovery. **f**, MitoScribe records a permanent record of the transient hypoxia exposure. Circles show each of the 3 biological replicates. **g**, Hypoxia responsive MitoScribe writer applied to MDA-MB-231 breast cancer cells mimics heterogeneous hypoxia response across the population. **h**, Quantification of hypoxia recording in MDA-MB-231 cells that harbored a MitoScribe hypoxia writer exposed to normoxia (21%), mild hypoxia (8%), or strong hypoxia (2%). Circles and error bars show the mean ± s.d. of 3 independent biological replicates. P < 0.0001 by one-way ANOVA followed by Tukey post hoc test.

We next tested whether the recording could be used to recover a past history of hypoxia. We transiently exposed cells harboring the hypoxia MitoScribe writer to low oxygen for a single day, then transitioned the cells back to normoxia for continued growth (Fig. 2d). The hypoxic exposure led to increased transcription of the MitoScribe writer as well as the hypoxia-responsive genes VEGF and HIF1a(*41*), however, this transcriptional response to transient hypoxia returned rapidly to baseline and was entirely undetectable two days after recovery (Fig. 2e). In contrast, the MitoScribe recording increased following the hypoxic exposure, stabilized in normoxia, and remained at ∼4.5% as a permanent log of the prior exposure, which could be recovered 2 weeks later after many cell divisions and passages in culture (Fig. 2f).

To test the MitoScribe hypoxia recorder in more physiologically relevant cell type, we integrated the writer into MDA-MB-231 breast cancer cells using a piggyBac transposon system, and cultured these cells in varying levels of oxygen for 72 h to mimic the oxygen gradients in tumors (Fig. 2g). As expected, cells cultured at 2% O_2_ showed recording of the hypoxic exposure with 0.9% editing, whereas intermediate (8%) or normoxic (21%) conditions showed no significant recording at 0.008% and 0.016%, respectively (Fig. 2h). These results demonstrate that MitoScribe can durably encode environmental exposure into the mitochondrial genome with single-nucleotide precision.

### Recording of signal strength

In addition to capturing the exposure and temporal persistence of a biological signal, we next explored whether MitoScribe could be used to distinguish between graded inputs, which would enable the quantitative recording of signal strength. To address this, we focused on the Wnt pathway, a key regulator of development and stem cell fate. We first used a Wnt writer with a promoter containing 8 repeats of the TCF-LEF Wnt responsive element and activated the pathway using different concentrations of CHIR99021, a GSK3β inhibitor for 48 h. Wnt-recording levels increased in a dose-dependent manner from 0-10 µM, with higher dosage of CHIR inducing greater editing efficiency up to 19.8%, illustrating the capacity to distinguish between levels of a graded response (Fig. 3a,b). As an alternative way of creating a graded response, we used a single concentration of 5 µM CHIR99021, but varied the number of WRE repeats to create Wnt responsive promoters of different strengths. We found that decreasing the number of response elements led to a corresponding decrease in Wnt recording in the presence of a uniform CHIR dose for 48 h once the number of repeats was reduced below four, illustrating the capacity to record graded promoter strength (Fig. 3c,d).

**Figure 3.**
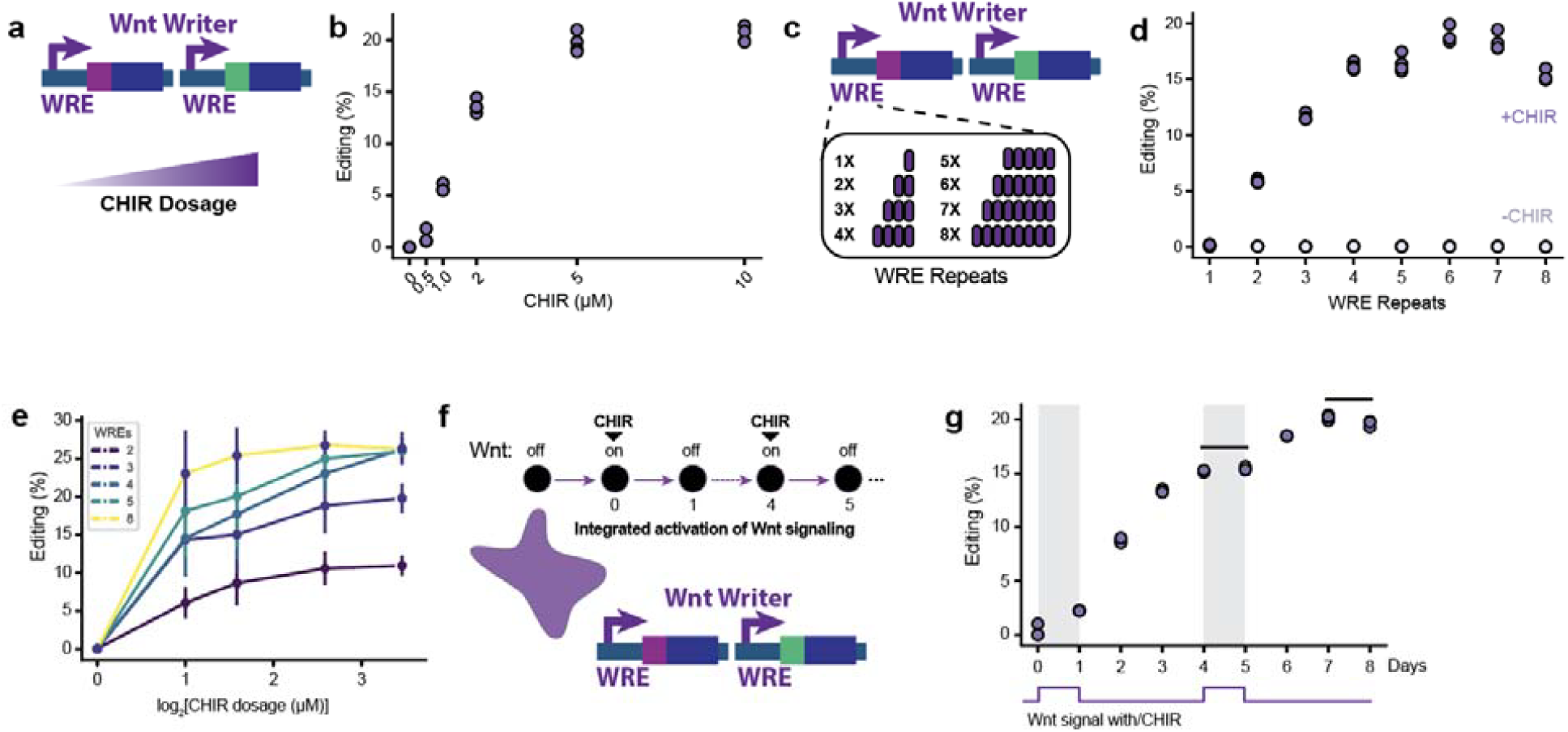
Recording of signal strength. **a**, Schematic of a dose-response treatment of HEK293T cells carrying Wnt signaling responsive writer. **b**, Increase in CHIR dosage from 0 to 10 µM is reflected in the editing percentage. Circles show each of 4 biological replicates. **c**, MitoScribe Wnt writer variations with different numbers of WRE repeats. **d**, The number of WREs from 1 to 8 is reflected in the editing percentage. Circles show each of 4 biological replicates. P < 0.0001 by two-way ANOVA followed by Sidak correction. **e**, Combinatorial matrix of CHIR dosage and WRE number shows editing scales with signal strength across a broad dynamic range. Circles and error bars show the mean of 3 biological replicates ± s.d.. **f**, Experimental workflow for integrated CHIR pulses of HEK293T cells harboring a MitoScribe Wnt writer separated by washout periods. **g**, Quantification of editing over time shows stepwise increases with each 5 µM CHIR exposure pulse, showing that MitoScribe integrates multiple transient signaling events into cumulative, stable mitochondrial edits. Grey area represents the CHIR exposure pulse. Circles show each of the 3 biological replicates.

We next leveraged the differences in Wnt responsive promoter strengths to evaluate the tunability of the recording system over a broad range of Wnt signaling intensities. To do so, we varied both the CHIR dosage and the WRE copy number. The recording scaled with input strength; that is, MitoScribe writers with a higher number of WREs were more sensitive to small changes in CHIR concentration, but saturated at lower doses, whereas writes with fewer WREs were less sensitive, but functioned over a larger CHIR dose range (Fig. 3e).

Next, we asked whether MitoScribe could integrate a more complex pattern of Wnt activity, with pulses of CHIR exposure separated over time (Fig. 3f). We observed that a single day of CHIR exposure, followed by 3 days of removal, led to a stable recording level that plateaued at ∼15% after the initial pulse. A second round of CHIR exposure led to an increase in recording and a new plateau of up to ∼20%, indicating that MitoScribe accumulates records in a stepwise manner and reflects the cumulative history of pathway activation (Fig. 3g). Taken together, our results demonstrate that MitoScribe captures both the intensity and duration of signal pathway activation in a cumulative and tunable manner.

### Multi-event recording

Having established that MitoScribe can encode individual cellular events at single mtDNA locus, we next extended its capacity to record multiple events simultaneously within the same cell. To do this, we designed multiple sets of writers driven by different promoters, each making neutral edits to adjacent regions of mtDNA. We selected 2 cytosines within the *MT-ND5* gene that generate synonymous mutations when transitioned to thymine, positioned at separate loci (Site 1 and 2), each of which could be independently targeted by a DdCBE mtDNA editor (Fig. 4a). Site 1 was assigned to a Wnt-responsive writer under the control of tandem Wnt response elements (WREs), while Site 2 was targeted by a BMP-responsive writer regulated by BMP response elements (BREs)(*42*) (Fig. 4b). All writer components were co-transfected into HEK293T cells. Twenty-four hours after transfection, we treated cells harboring these multiplexed MitoScribe writers with either CHIR or BMP. We found that treatment with 5 µM CHIR for 48 h, which activates Wnt signaling, led to 17.5% editing at Site 1 exclusively, with no editing at Site 2 (Fig. 4c). In contrast, stimulation of BMP signaling by its inducer, 200 ng/ml recombinant BMP2 or BMP4(*43*), induced editing of 6.4% or 5.6% at Site 2, respectively, with no editing at Site 1. Finally, combined stimulation with both CHIR and BMP ligands led to editing at both loci. This demonstrates that MitoScribe can record multiple signaling events independently within the same cells by targeting separate mtDNA sites with pathway-specific writers.

**Figure 4.**
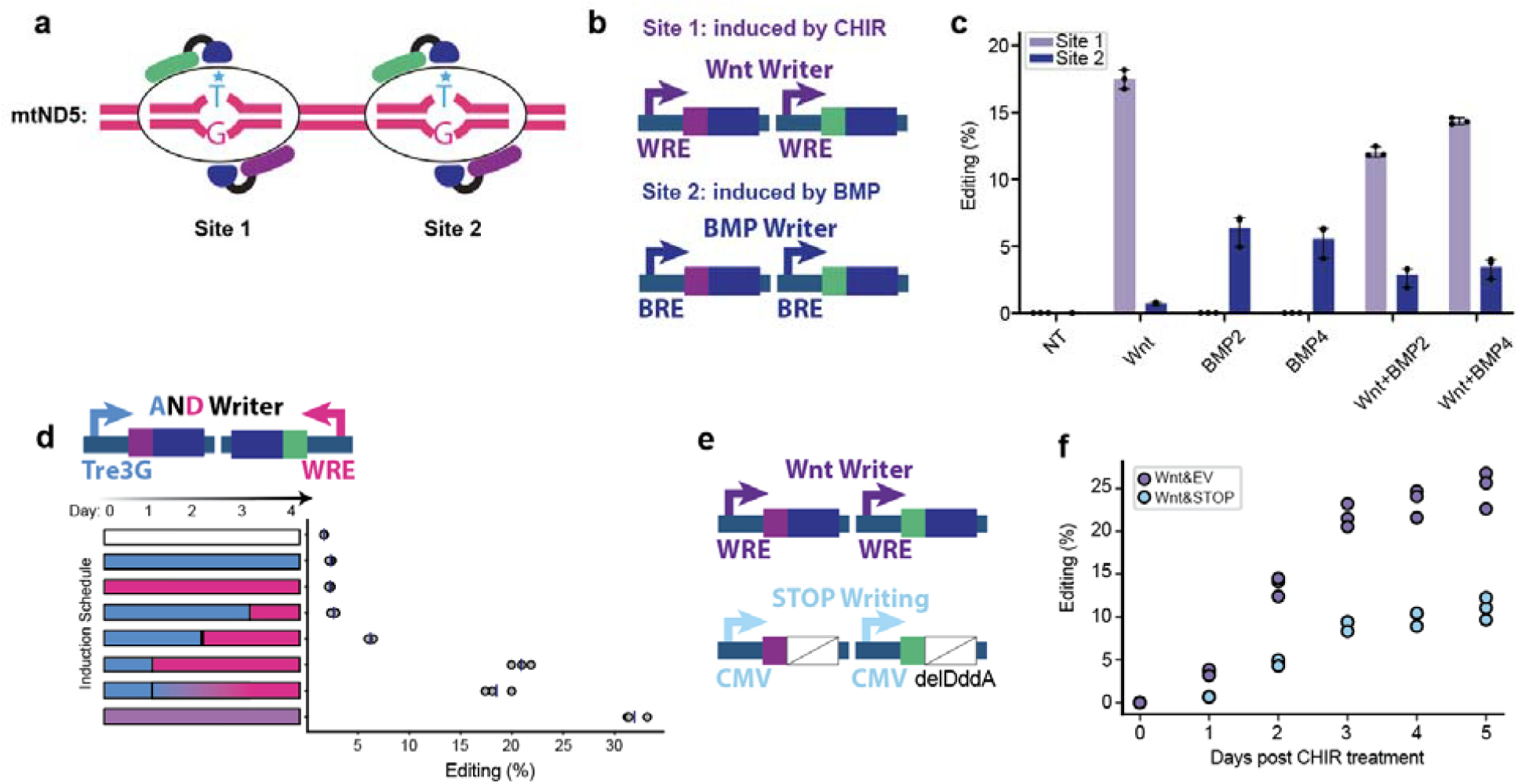
Multi-event recording and shutoff MitoScribe devices. **a**, Schematic of two distinct target sites within the *MT-ND5* gene, each carrying a separate synonymous editing site for event recording. **b**, Design of two pathway-specific writers: Wnt Writer with WRE-driven promoters and BMP Writer with BMP responsive elements (BREs), each controlling expression of writer halves targeted to different sites on the *MT-ND5* gene. **c**, Editing outcomes under different stimulation conditions for 48 h in HEK293T cells. Wnt stimulation (5 µM CHIR) induces editing at Site 1; BMP stimulation (200 ng/ml BMP2 or BMP4) induces editing at Site 2; combined Wnt and BMP stimulation edits both sites. Circles show each of 3 biological replicates. Bars are mean ± s.d.. P < 0.0001 by two-way ANOVA followed by Bonferroni correction. **d**, The AND gate writer configuration records coincident activation of Tet-on and Wnt signaling, shown with different degrees of temporal overlap between Dox (Tet-on) and CHIR (Wnt) treatment. Circles in show each of the 3 biological replicates, and vertical lines show the mean. P < 0.0001 by one-way ANOVA followed by Dunnett’s post hoc test. **e**, A second cassette with no deaminase limits the recording. **f**, Time-course of CHIR-induced editing in cells co-transfected with Wnt Writer and either empty vector or STOP construct. Circles show each of the 3 biological replicates. P < 0.0001 by Student’s unpaired two-tailed t-test.

### AND logic for coincidence detection and temporal gating

We next created a different version of a multiplexed MitoScribe writer using more advanced logic processing for coincidence detection. Specifically, we implemented an AND gate by splitting the recorder halves between two orthogonal promoters: one responsive to the Tet-on system (TRE3G) and the other responsive to Wnt signaling (WRE) (Fig. 4d). In this configuration, recording requires the temporally overlapping activation of both pathways to reconstitute the functional recorder. We performed time-resolved induction on varying inducer schedules and observed that the extent of editing depended on the duration of overlap between the two inputs, with maximal recording of 32% occurring only when both stimuli overlapped for an extended period (Fig. 4d). Thus, MitoScribe can be configured to detect coincident signaling events and convert them into a single-nucleotide edit in mtDNA.

We next tested an imposed STOP control over the editing. To do this, we introduced a dominant-negative STOP writer where we deleted the deaminase domain DddA to create a catalytically inactive writer (Fig. 4e). This STOP writer was designed to compete with the active writer for binding to the target site, thereby suppressing editing. When we co-transfected the Wnt writer along with the WntSTOP writer and compared it to the Wnt writer along with an empty vectorWnt, we observed that the STOP writer reduced recording efficiency from 25% to 11% in response to CHIR after 5 days (Fig. 4f). While this STOP writer did not fully suppress the recording, it demonstrates the potential to add layers of experimenter control to the MitoScribe system.

### Reconstruction of signal history in mesodermal differentiation

Wnt signaling plays a central role in early embryogenesis, stem cell differentiation, and lineage specification, with both the level and timing of signaling influencing downstream cell fate decisions(*44–47*). To examine how MitoScribe could be used to study developmental signaling in a defined biological context, we focused on the role of Wnt signaling during mesodermal differentiation of human induced pluripotent stem cells (iPSCs). First, we generated two iPSC lines with integrated MitoScribe Wnt writers: one in which each half of the editor was stably integrated into a single allele of the safe-harbor CLYBL locus, and the other in which a piggyBac transposon system was used for random multicopy integration (fig. S3a). Both lines showed increased editing with CHIR stimulation for 3 days, but the piggyBac line showed significantly higher editing rates compared to the CLYBL-targeted line (fig. S3b), consistent with differences in transgene copy number, so we opted to use the piggyBac line for downstream experiments.

We next subjected the iPSCs harboring the Wnt-responsive MitoScribe writer to a mesodermal directed differentiation protocol initiated by 5 µM CHIR for 1 day followed by washout for 2 days (Fig. 5a and fig. S3c). We collected single-cell RNA sequencing on 24,638 control and 19,976 CHIR-treated iPSCs, using a 10X Genomics platform. While this sequencing will include transcripts from the mitochondrial genome, the low depth of single-cell RNA sequencing risks diminishing the dynamic range of the MitoScribe recorder. Therefore, we additionally used targeted enrichment of MT-ND5 transcripts from cDNA libraries(*48*) to increase the sequencing coverage on the transcript that contains the recording (Fig. 5b).

**Figure 5.**
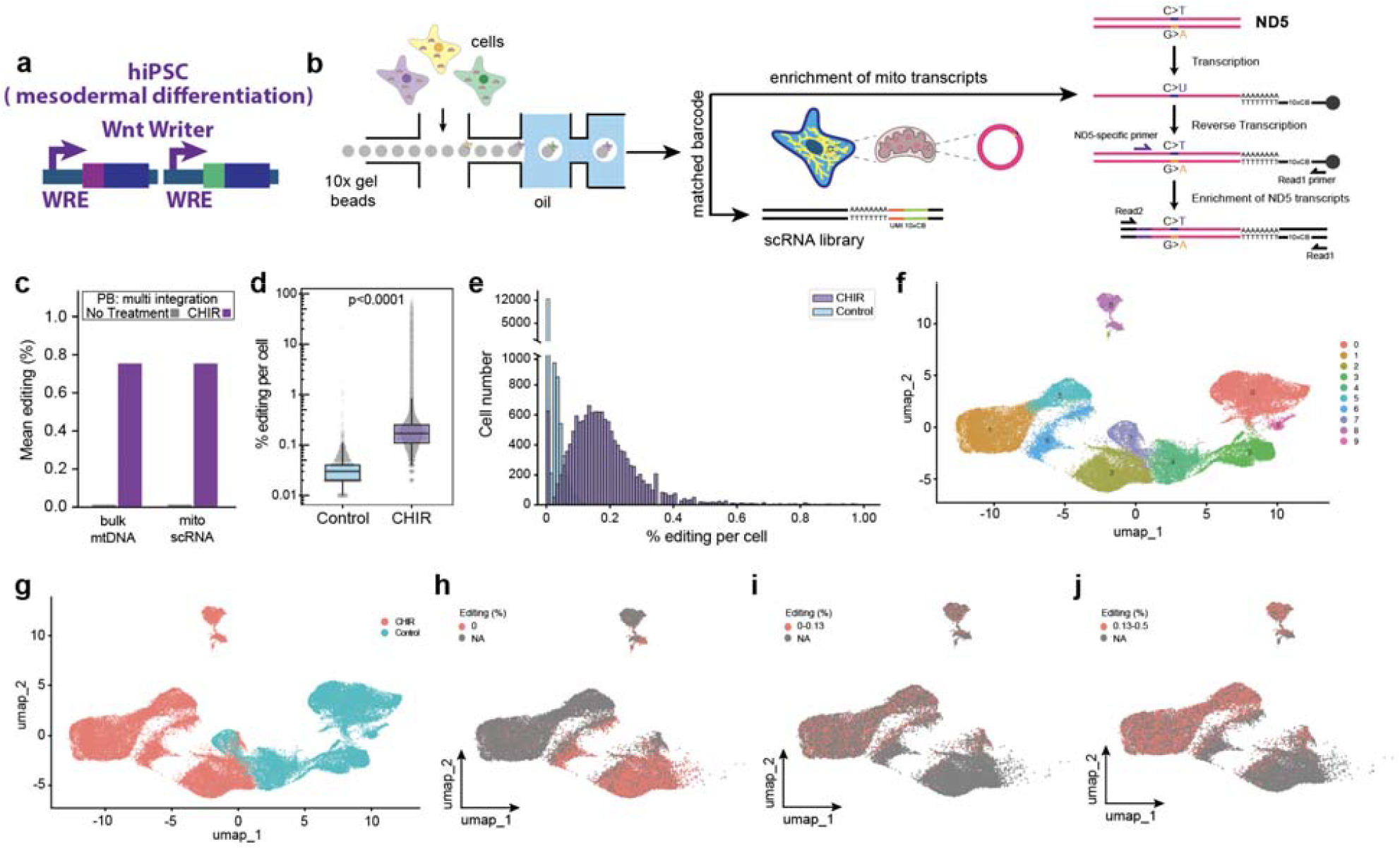
Reconstruction of signal history in early mesodermal differentiation. **a,** Wnt responsive MitoScribe writer in human induced pluripotent stem cells (hiPSC) during mesodermal directed differentiation. **b**. Experimental workflow of single-cell MitoScribe. Cells were encapsulated in microfluidic droplets for single-cell RNA sequencing (10x Genomics) with targeted enrichment of *MT-ND5* transcripts to quantify editing levels per cell alongside nuclear transcriptomes. Following mRNA capture and whole-transcriptome amplification, part of the cDNA is used for standard scRNA-seq, and another part is used for PCR-based enrichment of *MT-ND5* transcripts. The 150 bp sequencing reads maximize *MT-ND5 coverage fr*om the specific primer to the edits. **c**. Mean editing percentages measured in bulk mtDNA sequencing and single-cell mtRNA sequencing from piggyBac-integrated Wnt writer iPSCs, with and without 5 µM CHIR treatment. **d**. Single-cell box plot showing distribution of editing rates in control versus CHIR-treated iPSCs. P < 0.0001 by Student’s unpaired two-tailed t-test. **e**. Histogram showing the distribution of the number of cells by editing percentage per cell from 0-1% in the absence or presence of CHIR for Wnt stimulation, revealing continuous analog variation consistent with graded recording. **f**. UMAP projection of single-cell transcriptomes colored by identified clusters representing distinct mesodermal differentiation states. **g**, UMAP colored by treatment (CHIR versus control), showing separation of cell populations based on Wnt stimulation. **h**-**j**, UMAP overlays of editing rates per cell, binned into non (**h**), low (≤0.13) (**i**), and higher (>0.13–0.5) (**j**) categories, showing enrichment of highly edited cells in specific differentiation clusters.

For calling mtDNA variants, we separately sequenced the enriched MT-ND5 transcripts with 150-bp reads, then assigned each mtRNA to an individual cell using the cell barcode contained in Read1 and called the editing status of that RNA using Read2. We found that the editing rate that we obtained by bulk amplicon sequencing of this culture very closely matched the mean editing rate of all single cells determined by single-cell sequencing with mtRNA enrichment, demonstrating that the recording can be accurately reconstructed from single-cell RNA (Fig. 5c). After filtering for cells with >1,000 MT-ND5 reads, we identified 11,012 control and 11,749 CHIR-treated iPSCs with high-confidence depth for recording detection. Looking at the distribution of editing rate across single cells, we found that CHIR treatment resulted in higher editing than the untreated culture overall and, critically, that the individual cells in the CHIR condition fell across a distribution of editing values (Fig. 5d,e). This demonstrates the value of the single-cell MitoScribe approach to profile the analog range of responses to a common stimulus across a population of cells.

We next examined how the Wnt signaling response from the initial CHIR differentiation signal correlated with cell state at a later point in differentiation. Using uniform manifold approximation and projection (UMAP) for dimensionality reduction, we co-clustered the CHIR-treated and control cells based on mRNA expression data. We found multiple distinct clusters and clear separation between the CHIR-treated and control cells (Fig. 5f,g). Next, we mapped the single-cell editing rates for the CHIR-treated cells onto this UMAP. We found that cells with no editing despite having an mtRNA read depth >1000 reads were located largely in the bottom right clusters of the UMAP (Fig. 5h). Cells with editing percentages on the lower (left) side of the distribution of edited cells shown in Figure 5e (>0%, ≤0.13%) mapped to a non-overlapping region of the UMAP with the unedited cells, toward the left-side clusters (Fig. 5i). Finally, cells with editing percentages on the higher (right) side of the distribution of edited cells (>0.13%, ≤0.5%) were located in the top left clusters of the UMAP (Fig. 5j). The clusters associated with higher levels of editing (1, 5, 7, and 8) were positive for mesodermal marker genes including TBXT(*49*), and even cardiac mesodermal genes such as Hand1(*50, 51*), and GATA4(*52*), while being negative for the pluripotency marker Oct4(*53*) (Fig. S4a-d). Thus, the history of Wnt signaling in these cells, captured in the MitoScribe recording, shows a heterogeneous response across iPSCs to the earliest differentiation cue, which predicts the later cell state, with larger responses associated with more differentiated states.

## DISCUSSION

This work demonstrates a new approach to log a history of graded biological signals at single-cell resolution using neutral base edits to the mitochondrial genome. The approach is flexible, with the ability to record multiple different signals, including hypoxia, NF-κB activity, BMP and Wnt signaling shown here. It is stable, with recordings maintained for weeks across dozens of cell divisions. It is analog, able to discriminate between the strength or duration of a given signal. It is portable, with recordings demonstrated in three different human cell types. It is multiplexable, able to record two signals independently or the temporal overlap between two signals. Finally, it has single-cell resolution, leveraging the large population of mitochondrial genomes within each cell to record the extent of a signal in the percentage of mtDNA molecules edited. These single-cell recordings can be recovered from RNA along with single-cell transcriptomes, to yield both the current cell state and the cell history simultaneously.

To put this technology in context with other approaches for logging molecular events, it may first be useful to state explicitly what it is not. It is not a lineage recorder, although it may be interesting in future work to combine MitoScribe with existing technologies for recording cell lineage. There is a temporal component to the coincidence detection of the *AND* format, but MitoScribe is not intended for determining the temporal sequence or exact timing of molecular events, which has been the focus of other molecular recording technologies(*21, 24, 27, 30*). Like most, but not quite all(*31*), molecular recorders, MitoScribe recordings are focused on a defined biological signal and are not aimed at open ended discovery.

MitoScribe *is* an analog event recorder. Previous analog event recorders have been implemented in bacteria that rely on the accumulation of mutations, including two that have been quite influential on our work: SCRIBE(*12*), an early analog recorder that used retron recombineering to accumulate mutations in response to signals; and CAMERA(*19*), which used editing of plasmids to store records of graded signals. While both of these have mammalian extensions, mSCRIBE(*17*) and CAMERA 2, the mammalian versions target genomic DNA and thus lose the dynamic range to recover single-cell graded recordings, which puts them in a separate category, also including ENGRAM(*25*), of approaches that are analog, but only at the population level.

There are multiple analog molecular recorders that do achieve single cell resolution in mammalian cells using imaging as the readout. Two interesting and conceptually similar approaches use long protein filaments as the storage material rather than DNA(*54, 55*). These approaches are capable of recording graded signals in the accumulation of fluorescent proteins or tags that can be immunostained and imaged by fluorescence microscopy. Another recent approach, INSCRIBE, uses base edits to genomic arrays followed by *in situ* transcription, hybridization, and fluorescence microscopy to achieve analog single cell recordings also read out by imaging(*56*).

MitoScribe and the imaging-based analog recorders outlined above have complementary strengths. While both are capable of single-cell resolution, the imaging approaches link molecular histories of cells to their physical cellular context, including morphological features and tissue localization. Whereas, MitoScribe recordings link the molecular histories of cells to their current transcriptional state due to the ability to read them out along with deep single-cell transcriptome profiling. Future development may begin to see a blending of these strengths and potentially spatial readout of the MitoScribe signal, but, for now, the technology one would choose will be dictated by the use case.

In this work, we used the MitoScribe technology to log the response to a differentiation cue within individual cells and associated that response to a later differentiated cell state, showing that the early response to Wnt signaling predicts the resulting cell state. This result is consistent with a theory of developmental heterogeneity(*57–59*) where seemingly identical cells, in this case the iPSC population, are in fact fluctuating though states of non-genetic micro-heterogeneity due to transcriptional bursts, cell cycle asynchrony, and other cellular phenomena. A uniform input, in this case CHIR treatment, applied to a micro-heterogeneous population may drive differential responses across cells, yielding macro-heterogeneity in the resulting cell fates. While this mechanism for developmental heterogeneity has been long theorized(*60*), there is little direct experimental evidence to prove it due to the technical difficulty in relating macro-heterogeneous cell state back to a previous molecular signaling response. This is a major strength of the MitoScribe approach with many similar applications across developmental biology. Moreover, the utility can be extended to other fields where the cause (signal) and effect (cell state) are separated over time, including cancer progression, immunological responses, cellular degeneration, and more.

## METHODS

### Plasmid construction

All the plasmids and sequences of primers and signal-responsive elements (SREs) used in this paper are listed in Supplementary Table 1 and 2. Sequences encoding mitoDdCBE variants targeting *MT-ND5* (site 1 and site 2) and *MT-ND6*(*34*) were synthesized as gene blocks (Genscript) and cloned into pCMV based expression vector. The hypoxia response element, NF-κB response element, BMP, and TCF-LEF response element were obtained from the hypoxia(*40*), NF-κB(*38*), BMP(*42*), and TCF-LEF reporter(*37*), and sub-cloned into the promoter region of mitoDdCBE vectors as MitoScribe writers, respectively.

For plasmid integration, piggyBac-SRE-DdCBE-ND5 constructs were cloned into two steps. First, sequences of SRE-DdCBE-ND5 havles were amplified from pSRE-DdCBE-ND5 vectors using Q5 Hot Start High-Fidelity DNA Polymerase (NEB). Second, piggyBac-TRE3G-slowFT-puro (pSLS.200) and piggyBac-TRE3G-NG2(1-10)-blast (pRFF.008) were digested with SpeI and BglII (NEB). The PCR products and digested vectors were purified, and then assembled using Gibson assembly (NEB). For piggyBac-TRE3G-DdCBE-ND5 constructs, sequences of DdCBE-ND5 were amplified and sub-cloned into pSLS.200 and pRFF.008 digested by NheI and BglII (NEB) as previous described(*61*).

### Mammalian cell culture

All cells were cultured and maintained at 37 °C with 5% CO_2_. Antibiotics were not used for cell culture of HEK293T cells, MDA-MB-231 cells and human iPSCs. HEK293T cells (CRL-3216, American Type Culture Collection (ATCC)) and MDA-MB-231 cells (a gift from Dr. Hani Goodarzi) were cultured in DMEM with GlutaMax (Thermo Fisher Scientific, 10566016) supplemented with 10% (v/v) fetal bovine serum (Gibco). WTC11 iPSCs (a gift from Dr. Bruce Conklin) were cultured on matrigel-coated T25 flasks (Corning, 430639) at 37 °C, 85% humidity, and 5% CO_2_. iPSCs were fed with mTeSR Plus medium (StemCell Tech, 100-0276) every other day. Upon reaching 80% confluence, iPSCs were passaged 1:5 and treated with 10 μM ROCK inhibitor (Y-27632 dihydrochloride, StemCell Tech, 72302). For maintenance, ReLeSR (StemCell Tech, 100-0483) was used to passage iPSCs as small colonies. For freezing cells and seeding specific numbers of cells for experiments, Accutase (StemCell Tech, 07920) was used to dissociate iPSCs as single cells after counting. All cell lines were tested negative for mycoplasma.

### Lipofection and nucleofection

Transfection of HEK293T and MDA-MB-231 was performed using Lipofectamine 3000 transfection protocol (Thermo Fisher Scientific, L3000015). Cells were seeded on 6-well TC-treated plates (VWR, 10861) at a density of 5 × 10_5_ cells per ml, 24 h before lipofection. Lipofection was performed at a cell density of approximately 70%. For MitoScribe experiments, cells were co-transfected with 1,200 ng of plasmids bearing mitoDdCBE halves to make up 2,400 ng of total plasmid DNA, and then were treated with signal agonists and collected at the indicated time points. For hypoxia induction, cells were cultured at 37 °C, 2% O_2_, and 5% CO_2_ and collected at indicated time points. For construction of MitoScribe line, cells were co-transfected with 1.6 µg of two cargo plasmids of piggyBac-SRE-DdCBE-ND5 halves and 800 ng of piggyBac transposase plasmid (pCMV-hyPBase) for piggyBac integrations. Stable cell lines were selected with 2 μg/ml puromycin (Gibco) and 10 μg/ml blasticidin S HCl (Gibco).

For nucleofection in human iPS cells and construction of MitoScribe iPSC line, 5 × 10^5^ cells were dissociated by Accutase and nucleofected with a plasmid mixture of 800 ng of piggyBac-WRE-DdCBE-ND5 halves and 400 ng of piggyBac transposase using the P3 Primary Cell Line 4D-Nucleofector X Kit (Lonza, program CA-137). After nucleofection, cells were seeded on 6-well matrigel-coated plates. The medium was changed after 24 h of nucleofection and cultured for 2 days before selection with 1 μg/ml puromycin and 5 μg/ml blasticidin S HCl. All the cell lines used in this paper are listed in Supplementary Table 3.

### Real-time quantitative PCR

Total RNA was extracted from HEK293T cells with the RNeasy Mini Kit (Qiagen). 100 ng of isolated RNA digested with RNase-free DNase set (Qiagen) was used for RT-qPCR. Analysis of hypoxia-downstream gene expression was performed with Luna Universal One-Step RT-qPCR (NEB, E3005S) using the primers listed in Supplementary Table 2. Data was normalized to β-actin abundance, and mRNA levels were calculated via the ΔΔCT method.

### Mitochondrial DNA level quantification

qPCRs were performed as previously described(*62*). Briefly, 10 ng of isolated DNA was used in the qPCR, performed with the use of the KAPA SYBR FAST qPCR Kit Master Mix (Roche Diagnostics) in a 10-μl reaction volume using the QuantStudio 5 Real-Time PCR System (Thermo Fisher Scientific). The relative abundance of the amplified *MT-ATP8* gene fragment was normalized to the amplified *β-actin* gene fragment. All the primers are listed in Supplementary Table 2.

### Western blot

For western blot analysis of MitoScribe writers expressed in mammalian cells, HEK293T cells were seeded on 6-well TC-treated plates (VWR) at a density of 4 × 10^5^ cells per ml, 18–24 h before lipofection. Cells were transfected with 1,200 ng of each mitoDdCBE half to make up 2,400 ng of total plasmid using Lipofectamine 3000. The turnover of proteins was analyzed by cycloheximide (CHX; Sigma-Aldrich, C7698) chase as previously described(*63*). Briefly, cells were treated with 100 μg/ml CHX and collected at the indicated time point. For preparation of cell lysate for western blot analysis of MitoScribe components, cells were harvested with a denaturing buffer containing 100 mM Tris-HCl, pH 8.0, and 1% SDS. Total cell lysates were sonicated and cleared by pelleting at 18,000g for 10 min. Protein concentrations were measured by BCA Protein Quantification Kit (Thermo Fisher Scientific, 23227). Next, 60 μl of cleared lysate supernatant was added to 20 μl of 4x LDS sample loading buffer (Thermo Fisher Scientific) with 1x Reducing Reagent (Thermo Fisher Scientific). Lysates were boiled for 10 min at 95 °C, and were used for gel electrophoresis, performed using Novex NuPAGE Bis Tris 4–12% gels (Thermo Fisher Scientific). Samples were separated by electrophoresis at 150 V for 60 min in Novex NuPAGE MES SDS running buffer (Thermo Fisher Scientific). Wet-transfer to a PVDF membrane was performed using the Mini Tran-Blot Cell (Bio-Rad) according to the manufacturer’s protocols. The membrane was blocked in 5% non-fat milk in TBST buffer for 1 h at room temperature, then incubated with Alexa 647-labelled mouse anti-DYKDDDDK (Thermo Fisher Scientific, MA1-142-A647; 1:1,000 dilution), Dylight 550-labelled mouse anti-HA (Thermo Fisher Scientific, 26183-D550; 1:1,000 dilution), and rabbit anti-GAPDH (LI-COR 926-42216; 1:2,000 dilution) in blocking buffer overnight at 4 °C. The membrane was washed three times with TBST (1x TBS and 0.1% (v/v) Tween-20, pH 7.4) for 10 min each at room temperature, then incubated with Dylight 650-labelled secondary antibody goat anti-rabbit (Thermo Fisher Scientific, 84-546) diluted 1:2,000 in blocking buffer for 1 h at room temperature, followed by washing three times for 5 min each with TBST, then imaged using ChemiDoc Imaging System (Bio-Rad).

### Signal recording with ligands and environment

Doxycycline Hydrochloride (Dox; Sigma, D3072) was purchased as 100 mg/ml stock in DMSO. Recombinant Human TNFI⍰ (PeProTech, 300-01A) was reconstituted in PBS as 10 μg/ml stock. CHIR99021 (StemCell Tech, 72054) was reconstituted in DMSO as 5 mM stock. Recombinant Human BMP2 (R&D, 355-BM) and BMP4 (R&D, 314-BP) was reconstituted in sterile 4 mM HCl (Sigma, H9892) containing 0.1% BSA (Sigma, A8412) as 200 μg/ml stork. All chemicals were aliquoted and stored at −20 °C and recombinant proteins were aliquoted and stored at −80 °C, thawed immediately before use and diluted with the appropriate culture medium. For dose-dependent recording experiments, HEK293T cells were seeded on a 12-well plate, 12 h post transfection with Wnt MitoScribe writers, and then were treated with various concentrations of CHIR99021 for 48 h. For the time-dependent recording experiments, HEK293T cells transfected with hypoxia MitoScribe writers were cultured in 2% O_2_-incubator and collected at various time points. For historical signal recording, HEK293T cells transfected with hypoxia MitoScribe writers were cultured in 2% O_2_-incubator for 24 h followed by culturing back to 21% O_2_-incubator to store hypoxia recordings, and harvested at indicated time points. For integrated signal recording, HEK293T cells integrated with Wnt MitoScribe writers were treated with 5 µM CHIR99021 for 24 h followed by washing with medium for 72 h, and then were performed the same treatment and washout as described. For AND gate recording, HEK293T cells integrated with AND gate writer (pLW.075 and pLW.088) were treated with 1 µg/ml Dox or 5 µM CHIR99021 at the indicated time points. Medium was changed every day for refreshing with the ligands or removal of the ligands. Both Dox and CHIR99021 were added of temporal overlap for coincidence detection. For all treatments, the same volume of DMSO or PBS was added to the medium as a negative control.

### Mesoderm Differentiation

Mesoderm differentiation (fig. S3c) under standard control conditions was performed as previously described before(*64*). Briefly, upon 50–70% confluency of iPSCs integrated with Wnt MitoScribe writers were seeded on matrigel-coated T75 flasks (Corning) and treated with 10 μM ROCK inhibitor (Y-27632) for 24 h. To start induction, Y-27632 was removed and 5 μM CHIR99021 was added in RPMI 1640 (Thermo Fisher Scientific, 61-870-036) supplemented with B27 minus insulin (Thermo Fisher Scientific, A1895601) for 24 h and removed with the medium exchange from 24 h to 72 h. Mesodermal cells were harvested for bulk mtDNA sequencing and sc-mtRNA sequencing 72 h post activation. Upon treatment, the same volume of DMSO was added to the medium as a negative control.

### High-throughput amplicon sequencing of DNA samples

Genomic DNA was extracted using a QIAamp DNA mini kit (Qiagen, 51306) according to the manufacturer’s instructions. DNA was eluted in 200IZµl of ultra-pure, nuclease-free water. Then, DNA yield was measured and normalized to 50 ng/µl. 100 ng of the gDNA was used as template in 25-µl PCR reactions with primer pairs to amplify the locus of interest, which also contained adapters for Illumina sequencing preparation (Supplementary Table 3). The amplicons were purified using AMPure XP beads (Beckman Coulter) according to the manufacturer’s instructions, and the amplicons were eluted in 20IZµl of ultra-pure, nuclease-free water. Finally, the amplicons were indexed and sequenced on an Illumina NextSeq instrument, with at least 50,000 reads per sample. FASTQ files were demultiplexed with bcl2fastq (Illumina, v.2.20) and processed with CRISPResso2(*65*) to quantify the percentage of on-target mtDNA edits.

### Single-cell RNA sequencing and analysis

Following dissociation of ∼20,000 WTC11 iPSCs and iPSC-derived mesodermal cells, the scRNA-seq library was prepared using Chromium Next GEM Single Cell 3IZ Reagent Kits v4 according to the manufacturer’s protocol (10X Genomics). The generated scRNA-seq libraries were sequenced using NovaSeq X aiming for 40K read pairs per cell. As previously described(*66*), raw sequencing data were aligned to the human reference genome (hg38) and preprocessed with the Cell Ranger v.9.0.0 pipeline (10X Genomics). Data from all samples was merged by CellRanger-aggr (10X Genomics) in the pipeline described above and normalized to the same sequencing depth, resulting in a single gene-barcode matrix. Further analysis was performed using Seurat v5.3.0 R package with reference to the Seurat web tutorials. Low quality cells were removed from analyses, keeping the cells with number of genes per cells between 2,800 and 10,000, with a UMI count per cell between 8,000 and 50,000, with mitochondrial gene percentage between 1 and 6%, and with ribosomal gene percentage between 10% and 25%. After the filtering step, for normalization, the data were transformed using SCTransform function, setting mitochondria gene percentages and ribosomal gene percentages and cell cycle scores as a variance to be regressed. Then, principal component analysis (PCA) was performed with RunPCA. Cells were clustered using top 30 principal components and visualized using a Uniform Manifold Approximation and Projection (UMAP) dimensionality reduction (RunUMAP, FindNeighbors, and FindClusters). For clustering, a vector of resolution parameters was passed to the FindClusters function and the optimal resolution that established discernible clusters with distinct marker gene expression was selected. To identify marker genes, the clusters were compared pairwise for differential gene expression using the FindAllMarkers function (min.pct = 0.25, logfc.threshold=1).

### Mitochondrial transcripts enrichment and data analysis

Targeted enrichment of mitochondrial transcripts was performed on single-cell cDNA libraries generated by the 10x Genomics 3’ v4 platform. PCR primers specific to the *MT-ND5* gene region were designed and modified following MAESTER protocol(*67*) to amplify this region while preserving cell barcodes and unique molecular identifiers (UMIs). The enrichment PCR was optimized to ensure high specificity and yield of *MT-ND5* transcripts from the whole-transcriptome amplified cDNA. The mtRNA enrichment consists of three PCR reactions with an AMPure XP beads clean-up in between. In brief, in PCR 1, 80 ng of the cDNA libraries were linearly amplified by 0.2 µM of only the reverse primer specifically target to *MT-ND5* in 25-µl PCR reactions (ten cycles of 98⍰°C for 15IZs, 62⍰°C for 30⍰s, and 72⍰°C for 1⍰min). PCR 2 serves to add adapters for Illumina sequencing preparation to mitochondrial transcripts while retaining the UMI and CB of the transcripts in 25-µl PCR reactions (twelve cycles of 98⍰°C for 15⍰s, 62⍰°C for 30⍰s, and 72⍰°C for 1⍰min). The third PCR is used to append Illumina adapters (P5 and P7), dual index barcodes to identify the sample, and sequencing primer binding sites in transcripts in 25-µl PCR reactions (six cycles of 98⍰°C for 15⍰s, 62⍰°C for 30⍰s, and 72⍰°C for 1⍰min). Final PCR products were purified using 0.7x SPRI beads to retain fragments approximately 400–1,200 bp in length and eluted in 20 µl of ultra-pure, nuclease-free water. The enriched libraries were sequenced on the Illumina NextSeq 2000 P2 platform with 28 cycles for Read 1, 150 cycles for Read 2, and 2 index barcodes. All primers used for enrichment of mitochondrial transcripts are list in Supplementary Table 3).

Raw sequencing reads were filtered for quality and aligned directly to the mitochondrial reference genome. Reads were grouped by cell barcode, UMI, and alignment position to collapse PCR duplicates into unique molecules, representing distinct original transcripts. Only reads with mapping quality (MAPQ) ≥ 30 and base quality (BQ) ≥ 30 were retained. Single-cell analyses included only cells with at least 1,000 collapsed molecules at the ND5 site to ensure statistical power. At the ND5 target site, the counts of molecules supporting the reference (C) and edited (T) reads were quantified per cell. The editing rate per cell was calculated by dividing the counts of edited (T) reads over the total counts of reference (C) reads and edited (T) reads.

Single-cell editing variant fractions were merged with matched 10x scRNA-seq metadata via cell barcodes. Data integration and visualization were conducted using the Seurat v5.3.0 R package, with the editing rate added as a metadata feature. The distribution of C to T substitution in the *MT-ND5* across cells was visualized on the existing UMAP embedding using Seurat’s FeaturePlot function.

## ACKNOWLEDGEMENTS

This work was supported by funding from the Bachrach Family Foundation, the Morgenthaler Family Foundation, and the National Institute on Aging (R01AG092732). S.L.S. is a Chan Zuckerberg Biohub – San Francisco Investigator. L.W. and N.P. were supported by a Fellowship from the California Institute for Regenerative Medicine (CIRM). We would like to thank Angelo Pelonero for his support on bioinformatic analysis methods.

## AUTHOR CONTRIBUTIONS

L.W.: Conceptualization, Methodology, Investigation, Formal Analysis, Visualization, Writing – Original Draft. N.P.: Investigation, Formal Analysis, Writing – Review & Editing. D.S.: Supervision, Writing – Review & Editing. S.L.S.: Conceptualization, Methodology, Supervision, Writing – Review & Editing, Project administration, Funding acquisition.

## DATA AVAILABILITY

Sequencing data associated with this study are available in the NCBI SRA (Bioproject PRJNA1311601)

http://www.ncbi.nlm.nih.gov/bioproject/1311601

## Supplementary Figures

**Figure S1.**
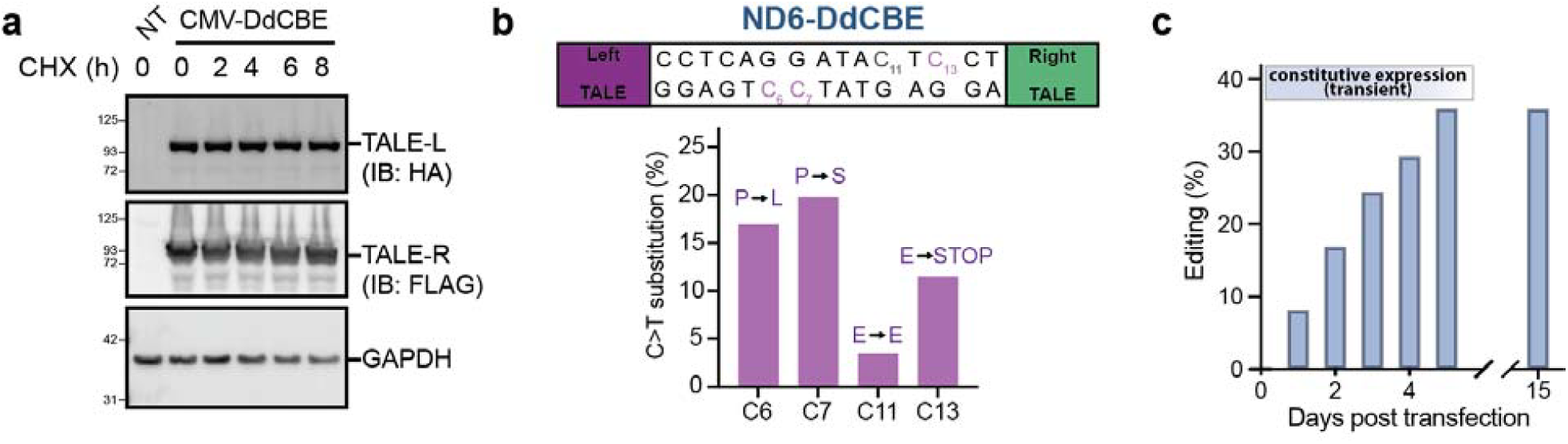
A synonymous mitochondrial edit enables durable and trackable recording. **a**, Western blot of constitutively expressed ND5-targeting DdCBE writer in HEK293T cells, showing expression and stability of HA-tagged writer left half and FLAG-tagged writer right half 2 days post-transfection. Cells were treated with 100 µg/ml cycloheximide (CHX) for the indicated times to monitor protein turnover. GAPDH was used as a loading control. **b**, The intended edit of the ND6-DdCBE and the frequency of missense, silent, and nonsense C>T substitutions. **c**, Time course of C>T substitutions at a synonymous site in the ND5 gene, introduced by MitoScribe under constitutive expression.

**Figure S2.**
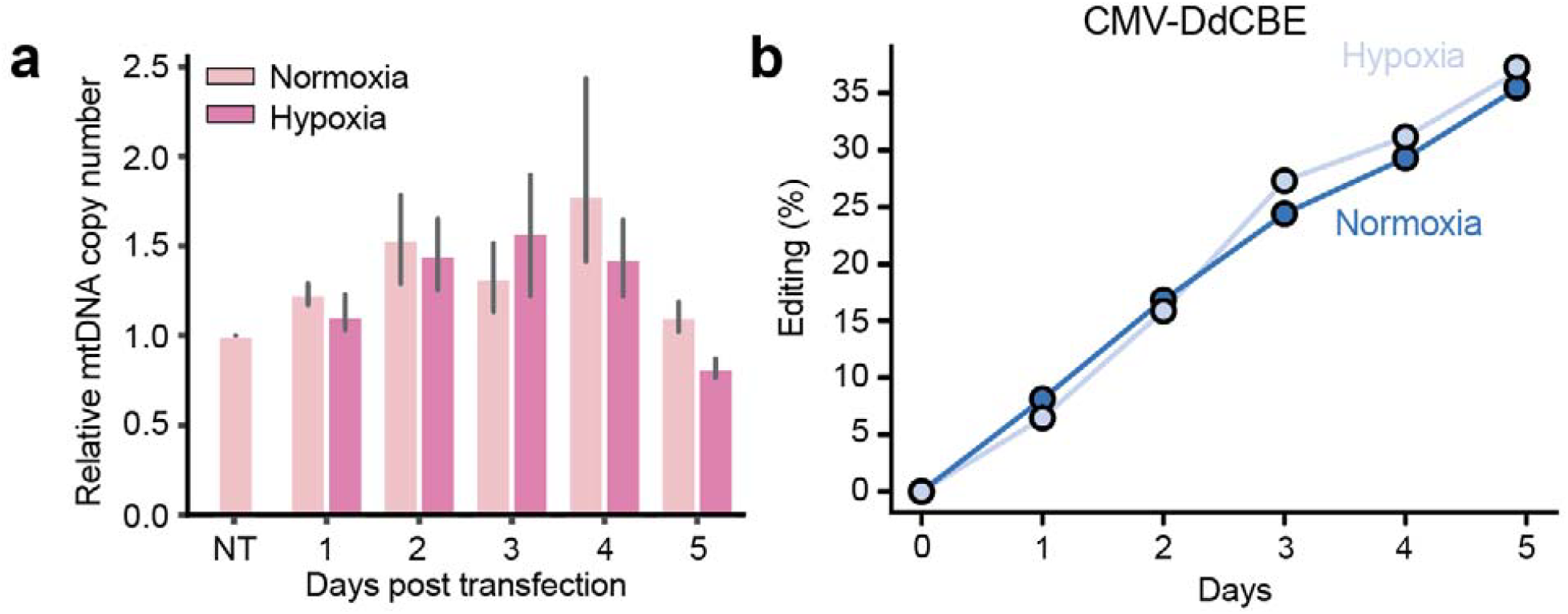
Hypoxia does not affect mitochondrial DNA copy number or the efficiency of constitutive DdCBE editing. **a**, Quantification of mitochondrial DNA (mtDNA) copy number in HEK293T cells transfected with hypoxia writer for 2 days in normoxic (21% O_2_, light pink) or hypoxic (2% O_2_, dark pink). mtDNA levels were measured by qPCR targeting the mitochondrial gene ATG8, normalized to β-actin, and presented relative to the non-transfected (NT) control. Bars represent mean ± s.e.m. of 3 independent biological replicates. **b**, Time-course experiment showing progressive accumulation of editing in HEK293T cells harboring the of constitutively expressed ND5-targeting DdCBE writer during 5 days of hypoxia and normoxia

**Figure S3.**
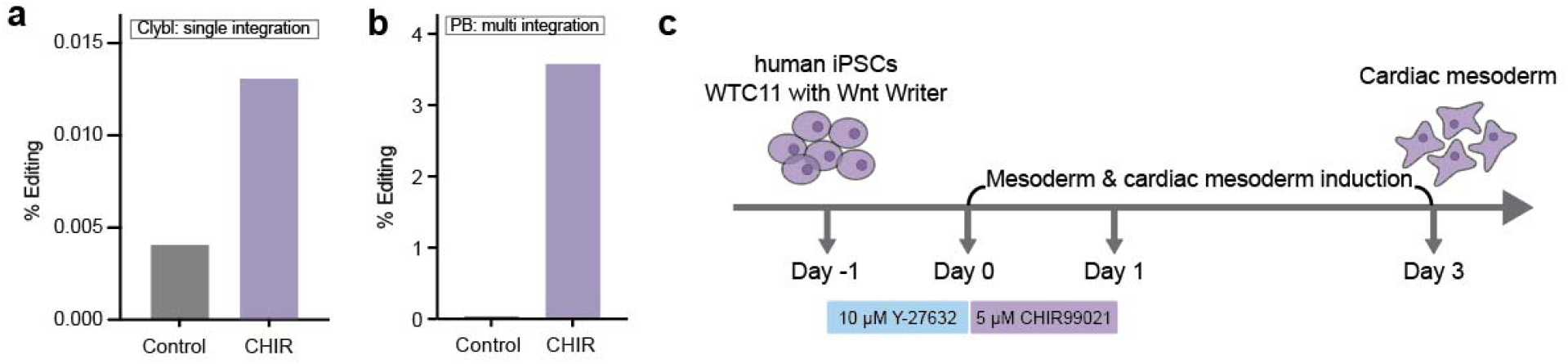
MitoScribe enables recording Wnt signaling in hiPSCs. **a**, Editing percentages of hiPSCs integrated with Wnt-responsive MitoScribe constructs into the CLYBL safe harbor locus, showing increased signal upon 5 µM CHIR99021 treatment for 3 days. **b**, Editing percentages of hiPSCs integrated with Wnt-responsive MitoScribe constructs by piggyBac transposon, showing increased signal upon 5 µM CHIR99021 treatment 3 days. **c,** Schematic of the cardiac mesoderm differentiation protocol from human iPSCs. Cells cultured in 10 µM ROCK inhibitor (Y-27632) were additionally treated with 5 µM CHIR99021 for 1 day followed by washout for 2 days. Controls remained untreated. Cells were collected on day 3.

**Figure S4.**
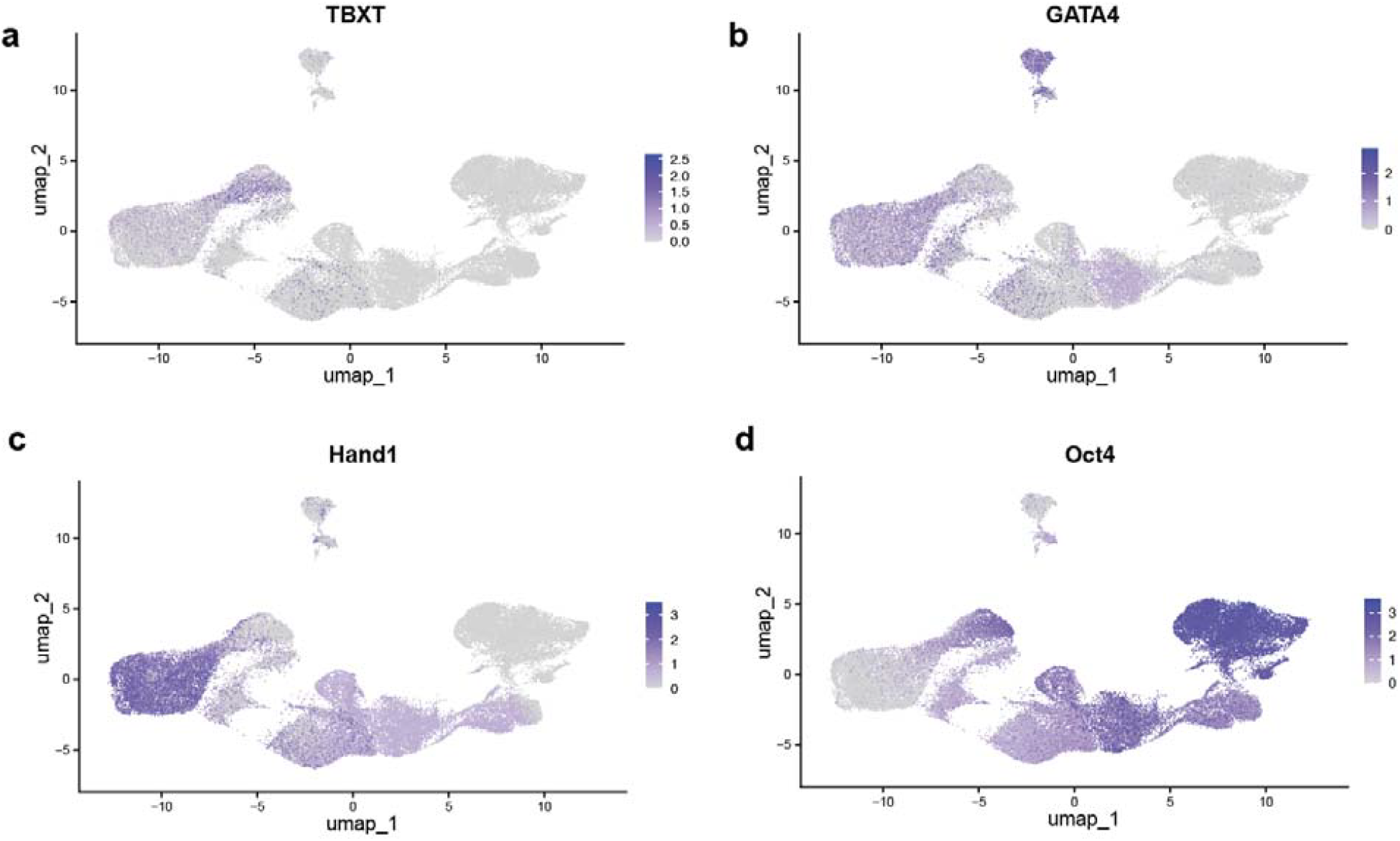
Single-cell transcriptomic validation of cardiac mesoderm induction in Wnt Writer–integrated hiPSCs. UMAP projection of single-cell transcriptomes, showing representative cardiac mesoderm markers of TBXT (**a**), GATA4 (**b**), and Hand1 (**c**), and the pluripotency marker Oct4 (**d**), derived from scRNA-seq analysis of control and differentiated WTC11 hiPSCs carrying the Wnt Writer constructs (related to Fig. 5).

