## SUPPLEMENTARY FIGURES for "MitoScribe single-cell molecular recorder logs graded signaling dynamics into mitochondrial DNA"

**Supplementary Figures**

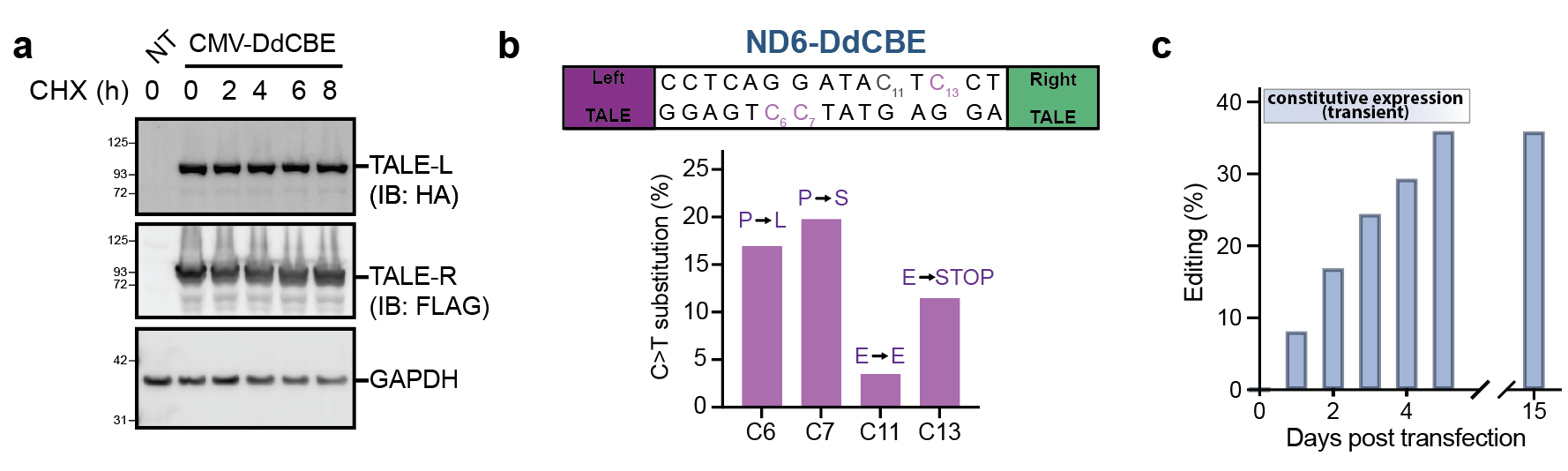

***Figure S1. A synonymous mitochondrial edit enables durable and trackable recording.***

**a**, Western blot of constitutively expressed ND5-targeting DdCBE writer in HEK293T cells, showing expression and stability of HA-tagged writer left half and FLAG-tagged writer right half 2 days post-transfection. Cells were treated with 100 µg/ml cycloheximide (CHX) for the indicated times to monitor protein turnover. GAPDH was used as a loading control. **b**, The intended edit of the ND6-DdCBE and the frequency of missense, silent, and nonsense C>T substitutions. **c**, Time course of C>T substitutions at a synonymous site in the ND5 gene, introduced by MitoScribe under constitutive expression.

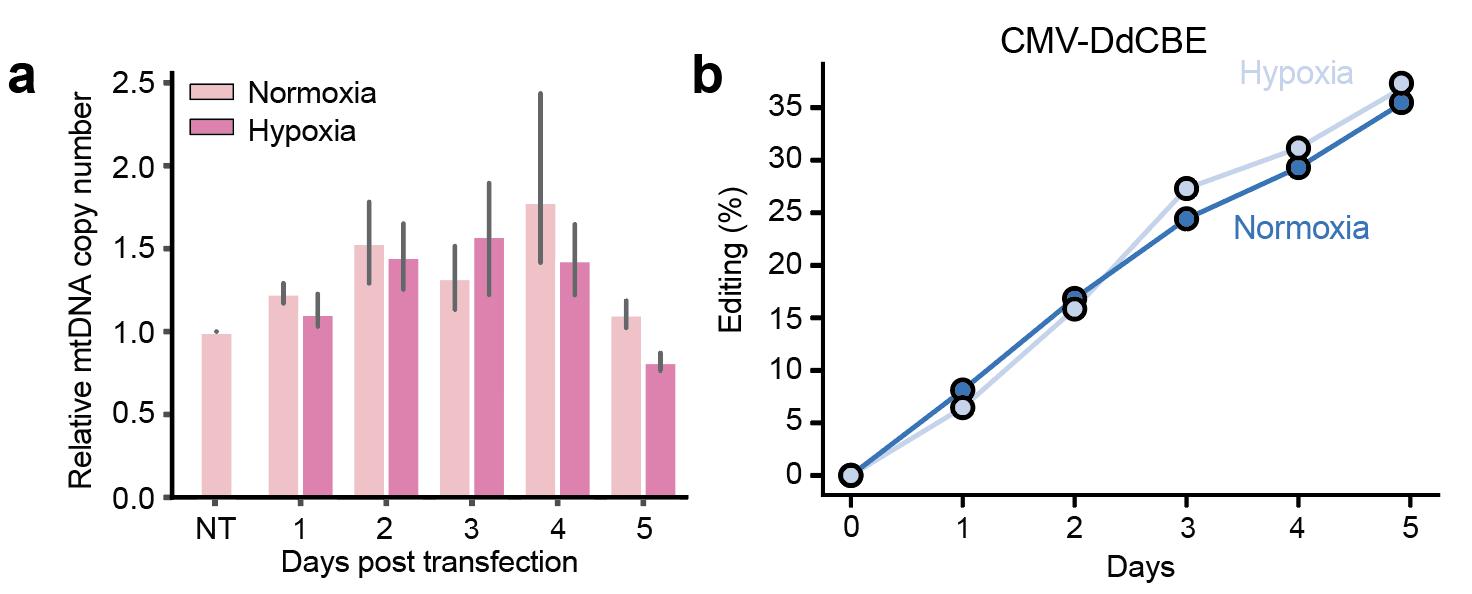

***Figure S2. Hypoxia does not affect mitochondrial DNA copy number or the efficiency of constitutive DdCBE editing.***

**a**, Quantification of mitochondrial DNA (mtDNA) copy number in HEK293T cells transfected with hypoxia writer for 2 days in normoxic (21% O₂, light pink) or hypoxic (2% O₂, dark pink). mtDNA levels were measured by qPCR targeting the mitochondrial gene ATG8, normalized to β-actin, and presented relative to the non-transfected (NT) control. Bars represent mean ± s.e.m. of 3 independent biological replicates. **b**, Time-course experiment showing progressive accumulation of editing in HEK293T cells harboring the of constitutively expressed ND5-targeting DdCBE writer during 5 days of hypoxia and normoxia
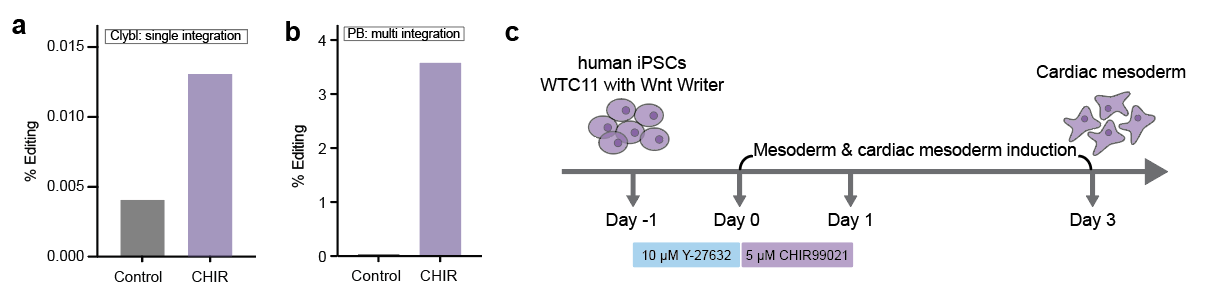

***Figure S3. MitoScribe enables recording Wnt signaling in hiPSCs.***

**a**, Editing percentages of hiPSCs integrated with Wnt-responsive MitoScribe constructs into the CLYBL safe harbor locus, showing increased signal upon 5 µM CHIR99021 treatment for 3 days. **b**, Editing percentages of hiPSCs integrated with Wnt-responsive MitoScribe constructs by piggyBac transposon, showing increased signal upon 5 µM CHIR99021 treatment 3 days. **c,** Schematic of the cardiac mesoderm differentiation protocol from human iPSCs. Cells cultured in 10 µM ROCK inhibitor (Y-27632) were additionally treated with 5 µM CHIR99021 for 1 day followed by washout for 2 days. Controls remained untreated. Cells were collected on day 3.

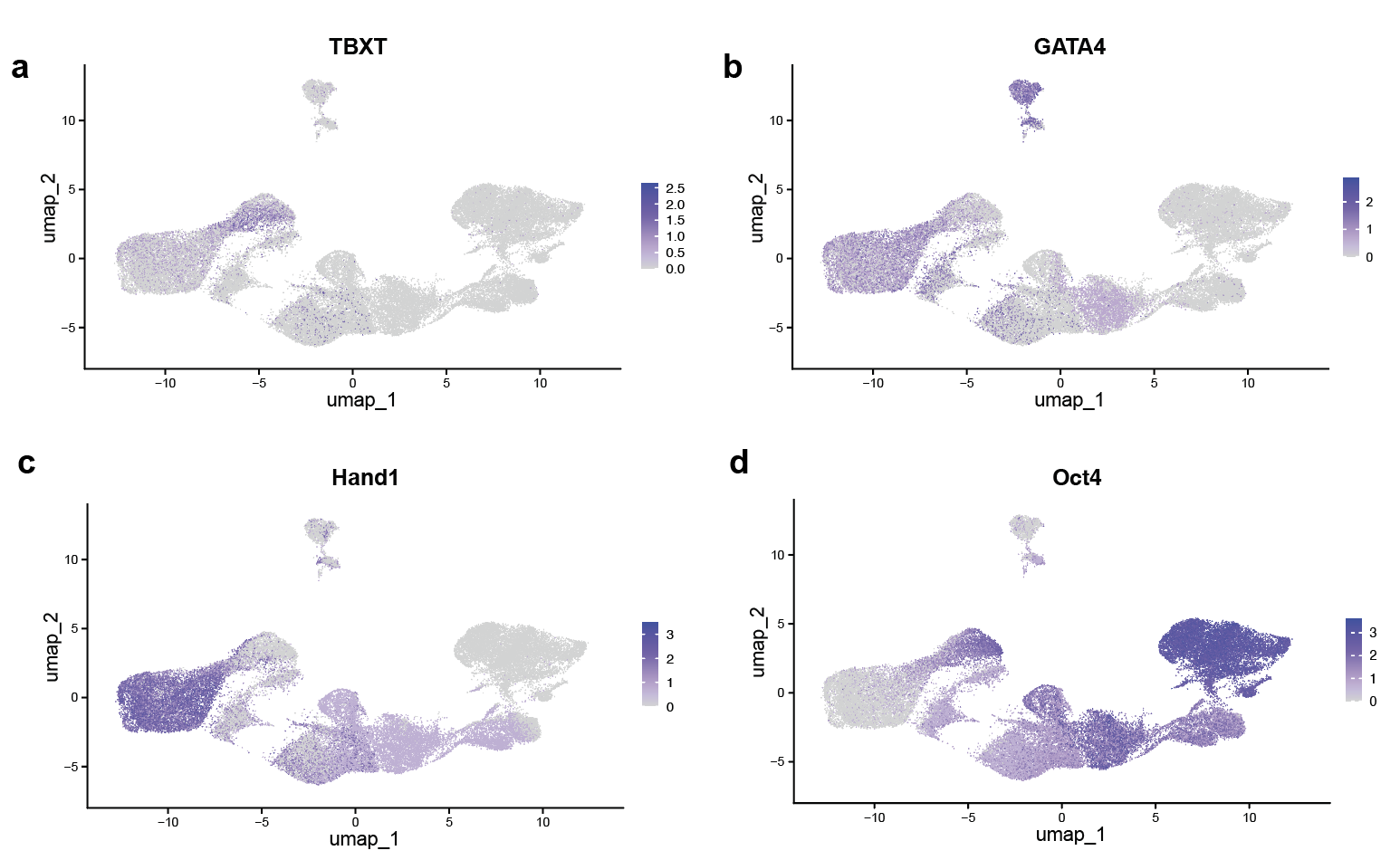

***Figure S4. Single-cell transcriptomic validation of cardiac mesoderm induction in Wnt Writer–integrated hiPSCs.***

UMAP projection of single-cell transcriptomes, showing representative cardiac mesoderm markers of TBXT (**a**), GATA4 (**b**), and Hand1 (**c**), and the pluripotency marker Oct4 (**d**), derived from scRNA-seq analysis of control and differentiated WTC11 hiPSCs carrying the Wnt Writer constructs (related to Fig. 5).
